# Knockout of the *SfVipR1* gene confers high-level resistance to *Bacillus thuringiensis* Vip3Aa toxin in *Spodoptera frugiperda*

**DOI:** 10.1101/2024.09.26.615236

**Authors:** Zheng Zhang, Lisi Wang, Xinru Pang, Wee Tek Tay, Karl H. J. Gordon, Tom K. Walsh, Yihua Yang, Yidong Wu

**Affiliations:** College of Plant Protection, Nanjing Agricultural University, Nanjing 210095, China; CSIRO, Black Mountain Laboratories, Clunies Ross Street, ACT 2601, Australia

**Keywords:** *Spodoptera frugiperda*, *Bacillus thuringiensis*, CRISPR/Cas9, Vip3Aa, resistance

## Abstract

**Background:** *Bacillus thuringiensis* (Bt) insecticidal proteins, including Cry proteins and vegetative insecticidal proteins (Vips), are extensively utilized in transgenic crops due to their efficacy and safety. The fall armyworm, *Spodoptera frugiperda*, has evolved practical resistance to Cry1Fa, yet no practical resistance to Vip3Aa has been documented. However, both laboratory selection and field screen studies indicate a high potential for this pest to evolve resistance to Vip3Aa, making it crucial to evaluate potential resistance genes. HaVipR1 has recently been identified as a key determinant of Vip3Aa resistance in the cotton bollworm, *Helicoverpa armigera*. This study investigated whether the *HaVipR1*-homologous gene in *S. frugiperda* (*SfVipR1*) is similarly involved in Vip3Aa resistance.

**Results:** We employed CRISPR/Cas9 technology to generate a homozygous knockout strain of *SfVipR1*. In comparison with the parent susceptible YJ-19 strain, the knockout strain (Sfru-KO) exhibited high-level resistance to Vip3Aa (>1875-fold) but showed no resistance to Cry1Fa. This acquired resistance to Vip3Aa is autosomal, recessive, and genetically linked to the deletion mutation in *SfVipR1* within the Sfru-KO strain of *S. frugiperda*.

**Conclusion:** Disruption of SfVipR1 results in high-level resistance to Vip3Aa, highlighting SfVipR1 has a critical role in Vip3Aa toxicity in *S. frugiperda*, despite the exact mechanism remaining unclear. Early detection of *SfVipR1* mutant alleles in the field is essential for developing adaptive resistance management strategies against *S. frugiperda*.

## 1 INTRODUCTION

*Bacillus thuringiensis* (Bt) is a gram-positive bacterium that can produce specific and potent insecticidal proteins, and Bt formulations have been successfully used as environmentally benign biopesticides for more than a century.^1^ Crops genetically engineered to produce insecticidal Bt proteins (Bt crops) have been commercialized since 1996, and the cumulative area of global planting of Bt crops from 1996-2019 exceeded one billion hectares.^2^ Due to long-term and extensive use of Bt crops, at least 26 cases of practical resistance to Bt crops have been documented in 11 pest species against 9 crystalline (Cry) Bt proteins.^3^ Resistance evolution by target pests is threatening the economic and environmental benefits of planting Bt crops.

Bt insecticidal proteins used widely in transgenic crops include Cry proteins, which are generated in the parasporal crystal during sporulation phase, and vegetative insecticidal proteins (Vips), which are secreted in the culture medium during vegetative growth phase of *B. thuringiensis*.^4,5^ Cry proteins do not share homology in sequence and binding sites with Vip proteins, so cross-resistance between them is none or almost none.^5–8^ Therefore, Vip proteins (mainly Vip3Aa) are commonly used in combination with Cry proteins (Cry1 and Cry2) in pyramided Bt crops to delay resistance evolution and expand the control spectrum.^9^ So far, 49 events of corn and 15 events of cotton producing Vip3Aa have been approved for commercial planting in several countries.^10^ Although no practical resistance to Vip proteins has been reported so far, early warnings of resistance to Vip3Aa have been documented in *Spodoptera frugiperda* in Brazil and *Helicoverpa zea* in the United States.^3,11,12^ In addition, laboratory strains with high-levels of Vip3Aa resistance have been established in at least six lepidopteran species, including *Helicoverpa armigera, Helicoverpa punctigera, H. zea, Heliothis virescens, S. frugiperda* and *Mythimna seprata*.^13–17^

Understanding the molecular mechanisms underlying Bt resistance can enhance our capacity to monitor, manage, and counter pest resistance to Bt proteins. Insights into the genetic mechanism of Bt resistance are largely derived from the analysis of resistance to Cry toxins. Such resistance is often linked to mutations in receptors, such as cadherins and ATP-binding cassette (ABC) transporters, which result in completely lost or reduced toxin binding to midgut epithelia in resistant larvae.^8,18^ Even though various lepidopteran strains have been established in the lab with resistance to Vip3Aa, the molecular mechanisms conferring resistance to Vip3Aa remain largely elusive.^8,19^ Only recently, three studies have indicated a relationship between genetic changes and Vip3Aa resistance in *S. frugiperda* and *H. armigera*. An association between the downregulation of a transcription factor gene (*SfMyb*) and a 206-fold resistance to Vip3Aa was identified in a laboratory-selected strain of *S. frugiperda*.^20^ Disruption of a chitin synthase gene (*SfCHS2*) was found to be associated with a 5560-fold resistance to Vip3Aa in another laboratory-selected strain of *S. frugiperda*.^21^ Meanwhile, a causative relationship between the disruption or reduced expression of the *HaVipR1* gene and high-level resistance to Vip3Aa (>1000-fold) was established in two field-derived strains of *H. armigera*.^22^

*S. frugiperda*, originally endemic to North and South America, has now invaded Africa, Asia, Europe, and Australia/Oceania, emerging as one of the most detrimental crop pests worldwide.^23^ Transgenic Bt corn, genetically engineered to express Cry1Fa, Vip3Aa or Cry1Fa+Vip3Aa, has been widely used to control *S. frugiperda* in several countries across the Americas.^24^ Following extensive selection by Bt corn, *S. frugiperda* has evolved practical resistance to Cry1Fa in Puerto Rico, Brazil, the mainland United States, and Argentina.^25–28^ Although no practical resistance has been documented to date for *S. frugiperda* to Vip3Aa,^3,24^ both laboratory selection and field screening studies suggest that there is a significant potential for this pest to develop resistance to Vip3Aa.^24^ Recently, HaVipR1 has been identified as a critical factor in Vip3Aa resistance in the cotton bollworm, *Helicoverpa armigera*.^22^ In the present study, we knocked out the *HaVipR1*-homologous gene (*SfVipR1*) in *S. frugiperda* with CRISPR/Cas9 technology, and found that disruption of *SfVipR1* conferred high-level resistance to Vip3Aa. Our results confirm that SfVipR1 is similarly involved in Vip3Aa resistance as HaVipR1 and that SfVipR1 is a critical determinant of Vip3Aa resistance in *S. frugiperda*.

## 2 MATERIALS AND METHODS

### 2.1 Insect strains and rearing

The YJ-19 strain of *S. frugiperda* was initially collected in Yuanjiang county, Yunnan Province of China in June 2019, and has been reared in the laboratory without exposure to any insecticides since its collection. Using the CRISPR/Cas9 gene editing system, a large fragment between exon 3 and exon 9 of *SfVipR1* gene in the YJ-19 strain was deleted. The created homozygous knockout strain of *S. frugiperda* was named Sfru-KO.

Larvae of *S. frugiperda* were fed on an artificial diet based on wheat germ and soybean powder at 26 ± 2°C, 60 ± 10% relative humidity, and a photoperiod of 14/10 h (light/dark). Adults were supplied with a 10% sugar solution.

### 2.2 Bt toxins and bioassays

Vip3Aa and Cry1Fa protoxins were purchased from the Institute of Plant Protection, the Chinese Academy of Agricultural Sciences (CAAS, Beijing, China).

Toxicity of Vip3Aa and Cry1Fa to *S. frugiperda* larvae was determined by diet overlay bioassays. Serial concentrations of toxin solutions were prepared by diluting the toxin stock suspensions with 0.01 M phosphate buffered saline (PBS, pH 7.4). Firstly, 1.2 ml of liquid artificial diet was dispensed into each well (surface area = 2 cm^2^) of 24-well plates. After the diet cooled and solidified at room temperature, 0.1 ml solution of Bt toxin was applied evenly onto the surface of the artificial diet. At the same time, the diet plates for control treatment were overlaid with 0.1 ml solution of PBS. After the overlaid solution dried, a single unfed neonate larvae that had hatched within 6 h was placed into each well of 24-well plates. Mortality was recorded after 7 days, larvae were considered dead if they were dead or still stayed in the first instar. PoloPlus software (LeOra Software, Berkeley, USA) was used to analyze the probit data and calculate the LC_50_ (the concentration of toxin killing 50% of tested larvae) and the 95% fiducial limits of the LC_50_.

### 2.3 Molecular cloning of SfVipR1

Total RNA was extracted from a pool of five midguts of the sixth instar *S. frugiperda* with Trizol reagent (Invitrogen, Carlsbad, CA, USA) according to the manufacturer’s instructions. First-strand cDNA was then synthesized with 1 μg of total RNA using the One-Step gDNA Removal and cDNA Synthesis SuperMix Kit (TransGen Biotech Co., Ltd., Beijing, China). Full-length cDNAs corresponding to *SfVipR1* was PCR amplified using specific primers (Table 1) with reaction mixtures containing 25 μl 2×Gflex PCR buffer (Takara Bio, Shiga, Japan), 2 μl each of the sense and antisense primers (10 μM), 2 μl cDNA, 1 μl Tks Gflex DNA polymerase (Takara Bio), and 18 μl ddH_2_O in a final volume of 50 μl. PCR conditions were 94 °C 3 min, 35 cycles of 98 °C 10 s, 55 °C 30 s, 72 °C for 1 min, and a final extension at 72 °C for 7 min. PCR products were separated on 1.25% agarose gels in Tris-acetate-EDTA buffer and stained with ethidium bromide. PCR products of the expected size were gel-purified with an AxyPrep Gel extraction kit (Axygen Biosciences, Union City, CA, USA), and then cloned into the pGEM-T vector (TransGen Biotech, Beijing, China). *Escherichia coli* Trans1-T1 cells (TransGen Biotech) were transformed with these constructs and the inserts sequenced by Sangon Biotech Co., Ltd. (Shanghai, China). A total of five clones were sequenced for the *S. frugiperda* YJ-19 strain.

**Table 1.**
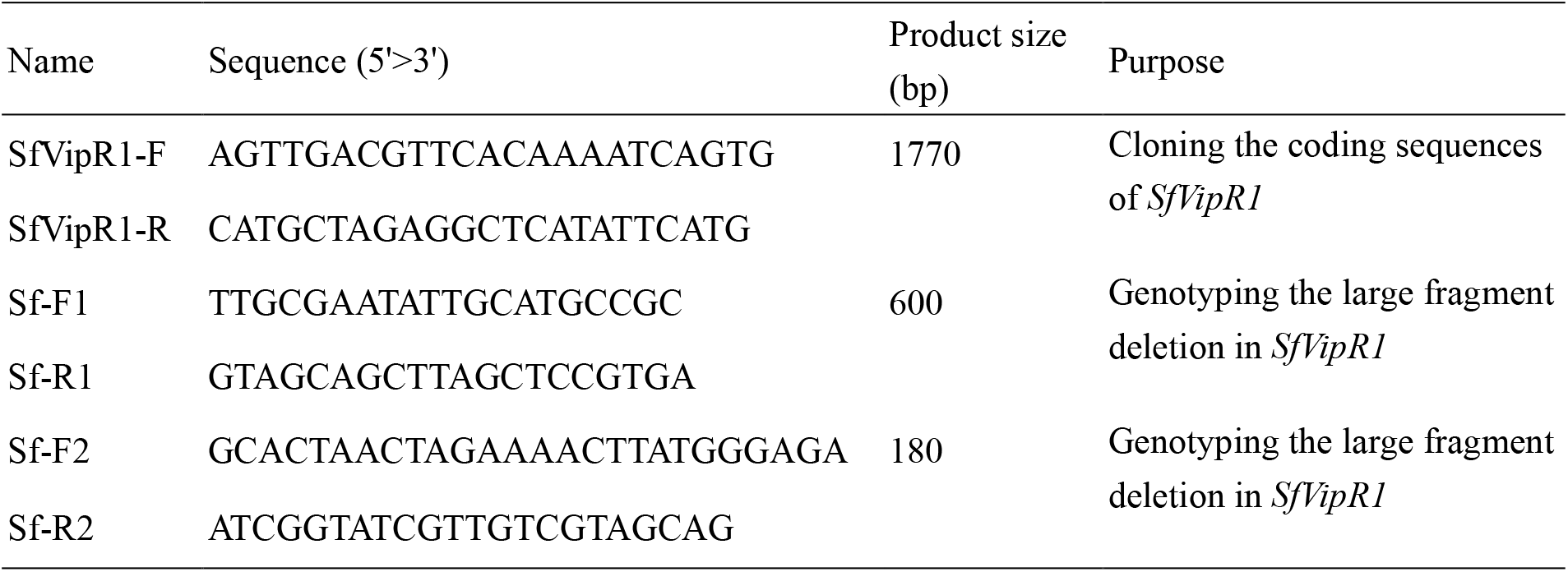
Primers used for genotyping the knockouts of *S. frugiperda*.

### 2.4 Design and preparation of sgRNAs

The sgRNA target sites of *SfVipR1* (GenBank no. PQ182785) were identified using the design principle of 5’-N_18-19_NGG-3’ (underlined is the PAM sequence) (Gene structure was sketched in Figures 2A). Two sgRNAs were used in this study (Sf-sgRNA1 targeting at exon 3 of *SfVipR1*: GGGCGACGGACTCTACTGTTGG; Sf-sgRNA2 targeting at exon 9 of *SfVipR1*: AGGAGGATCCAGTGTCGCCGAGG). Two oligonucleotides were used to synthesize sgRNA template: a forward primer harboring the T7 promoter sequence and the target sequence (5’-GAAATTAATACGACTCACTATAN_18-19_GTTTTAGAGCTAGAAATAGC-3’) and the universal oligonucleotide encoding the remaining sgRNA sequences (5’- AAAAGCACCGACTCGGTGCCACTTTTTCAAGTTGATAACGGACTAGCCTTATTTTA ACTTGCTATTTCTAGCTCTAAAAC-3’).

The fusion PCR reaction system (50 μl) consisted of 25 μl of 2 PrimeSTARMax Premix (TaKaRa, Dalian, China), 3 μl of 10 mM sgRNA-F, 3 μl of 10 mM sgRNA-R, and 19 μl of ddH_2_O ang performed at 98°C 30 s, 35 cycles of (98°C10 s, 60°C 30 s, 72°C 15 s), 72°C 10 min and hold at 12°C. The PCR products were purified using QIAprep® Spin Miniprep Kit (QIAGEN, Hilden, Germany). *In vitro* transcription of sgRNA was performed with MEGAshortscript™ T7 High Yield Transcription Kit (Ambion, Foster City, CA) according to the manufacturer’s instruction.

### 2.5 Cas9 protein and embryo microinjection

TrueCut™ Cas9 Protein v2 was purchased from Thermo Fisher (Shanghai, China). Fresh eggs (within 1 h after laid) were collected and soaked in 1% sodium hypochlorite solution for 2 min followed by washing with distilled water three times. After suction filtration, eggs were fixed on a glass slide with double-sides adhesive tape. Approximately one nanoliter mix of sgRNA1 (300 ng/μl), sgRNA2 (300 ng/μl) and Cas9 protein (200 ng/μl) were injected into individual eggs using a FemtoJet and InjectMan NI 2 microinjection system (Eppendorf, Hamburg, Germany). The whole process was completed in two hours. Injected eggs were placed at 26 ± 2°C with 60 ± 10% relative humidity and a photoperiod of 14/10 h (light/dark) for hatching.

### 2.6 Genomic DNA extraction and identification of mutations mediated by Cas9/sgRNA

Genomic DNAs of individual insects (moth legs or larvae) were extracted with AxyPrep™ DNA Extraction Kit (Axygen Biosciences, Union City, CA, USA) for genotyping purpose. Large fragment deletion mutations of individual *S. frugiperda* were determined by banding patterns of PCR fragments amplified with two pairs of specific primers of *SfVipR1* (Figure 2C). The direct sequencing of PCR products was conducted by TsingKe (Nanjing, Jiangsu, China) to detect the indel mutation types of *SfVipR1*. Sequences for the specific primer pairs were summarized in Table 1.

### 2.7 Inheritance of resistance to Vip3Aa

Fifteen pairs of adults were reciprocally mass crossed between the double knockout strain and the susceptible strain of *S. frugiperda*. Survival (%) of the knockout strain, the susceptible strain and their F_1_ progeny were determined at the diagnostic concentration of Vip3Aa (2 μg/cm^2^). According to the formula of Liu and Tabashnik,^29^ the dominance parameter *h* was calculated as: (survival rate of F_1_ - survival rate of susceptible strain) / (survival rate of knockout - survival rate of susceptible strain). The *h* values range from 0 (completely recessive) to 1 (completely dominant).

### 2.8 Genetic linkage analysis

Fifteen virgin female moths from the knockout strain were mass crossed with 15 male moths from the susceptible strain to produce F_1_. Fifteen male moths from the F_1_ progeny were backcrossed with 15 virgin female moths of the knockout strain to produce BC progeny. Neonates from the BC progeny were treated with the diagnostic concentration of Vip3Aa (2 μg/cm^2^) for 7 days using the bioassay methods described above. Individual survivors were randomly selected for DNA extraction and genotyping at the *VipR1* locus of *S. frugiperda*.

## 3 RESULTS

### 3.1 Characterization of the *S. frugiperda* VipR1 gene

The open reading frame sequence of *S. frugiperda VipR1* (*SfVipR1*) isolated from the YJ-19 strain contains 1275 nucleotides encoding a protein composed of 424 amino acids (GenBank no. PQ182785). Similarity of the deduced amino acid sequences is 55.26% between SfVipR1 and *H. armigera* VipR1 (HaVipR1) (see **Fig. 1**).

**Figure 1.**
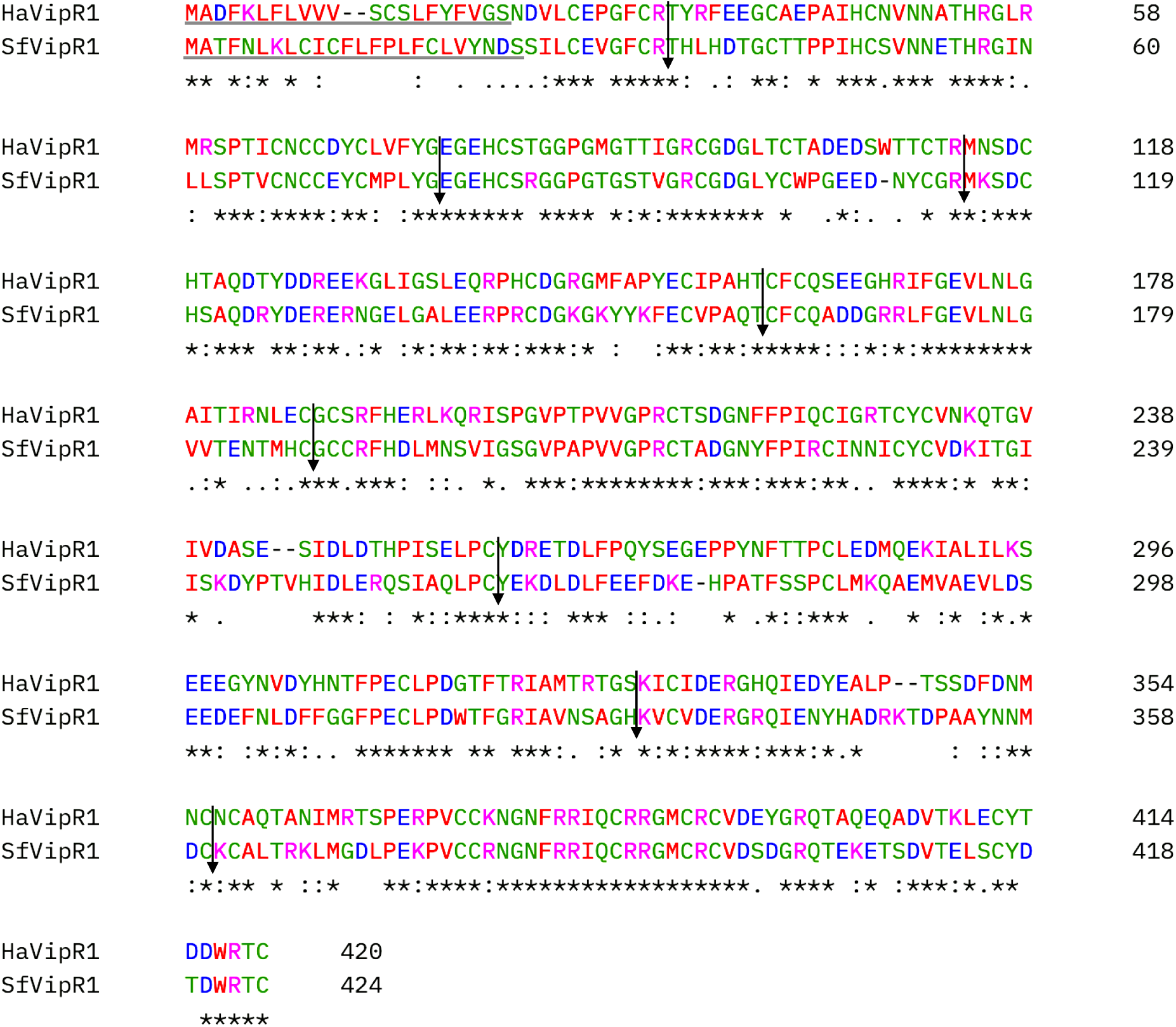
Alignment of VipR1 protein sequences with ClustalW for *Helicoverpa armigera* and *Spodoptera frugiperda*. Eight intron positions are marked with downward arrows. Signal peptide sequences underlined were predicted with SignalP 6.0 (https://services.healthtech.dtu.dk/service.php?SignalP). GenBank accession numbers are PZC80507.1 for *HaVipR1* and PQ182785 for *SfVipR1*.

Alignment of the full-length cDNA sequences with their respective genomic DNA sequences (NCBI Gene IDs: 110373801 for *HaVipR1*, 118279928 for *SfVipR1*) showed that the coding regions of *SfVipR1* and *HaVipR1* were interrupted by 8 introns at identical consensus amino acid positions, indicating that the genomic structure is conserved among the two *VipR1* genes (see **Fig. 1**). A signal peptide was predicted for SfVipR1 (24 aa) and HaVipR1 (21 aa), suggesting both proteins are targeted to the secretory pathway.

### 3.2 Construction of the SfVipR1 knockout strain (Sfru-KO) of *S. frugiperda*

Crossing and selection programs for establishing the homozygous *SfVipR1* knockout strain of *S. frugiperda* were shown schematically in **Fig. 2A**. 800 eggs of *S. frugiperda* YJ-19 strain were injected with a mixture of two sgRNAs and Cas9 protein. After injection, 230 eggs hatched. 73 G_0_ adults (35 ♂, 38 ♀) developed from the hatched eggs were crossed reciprocally with YJ-19 to produce G_1_. One leg from each of 150 G_1_ adults (pooled progeny of the reciprocal crosses) were cut out for genotyping to screen the individuals with a large fragment deletion in the *SfVipR1* gene. Six male and 7 female G_1_ adults heterozygous for a same large fragment deletion in *SfVipR1* were mass crossed to produce G_2_. Fifteen homozygotes (7 ♂, 8 ♀) with the large fragment deletion of *SfVipR1* identified from 70 G_2_ adults were pooled and mass crossed to establish the homozygous knockout strain, which is named Sfru-KO (**Fig. 3**). The large fragment deletion in *SfVipR1* results in knockout of almost the whole gene (seven out of the nine exons).

**Figure 2.**
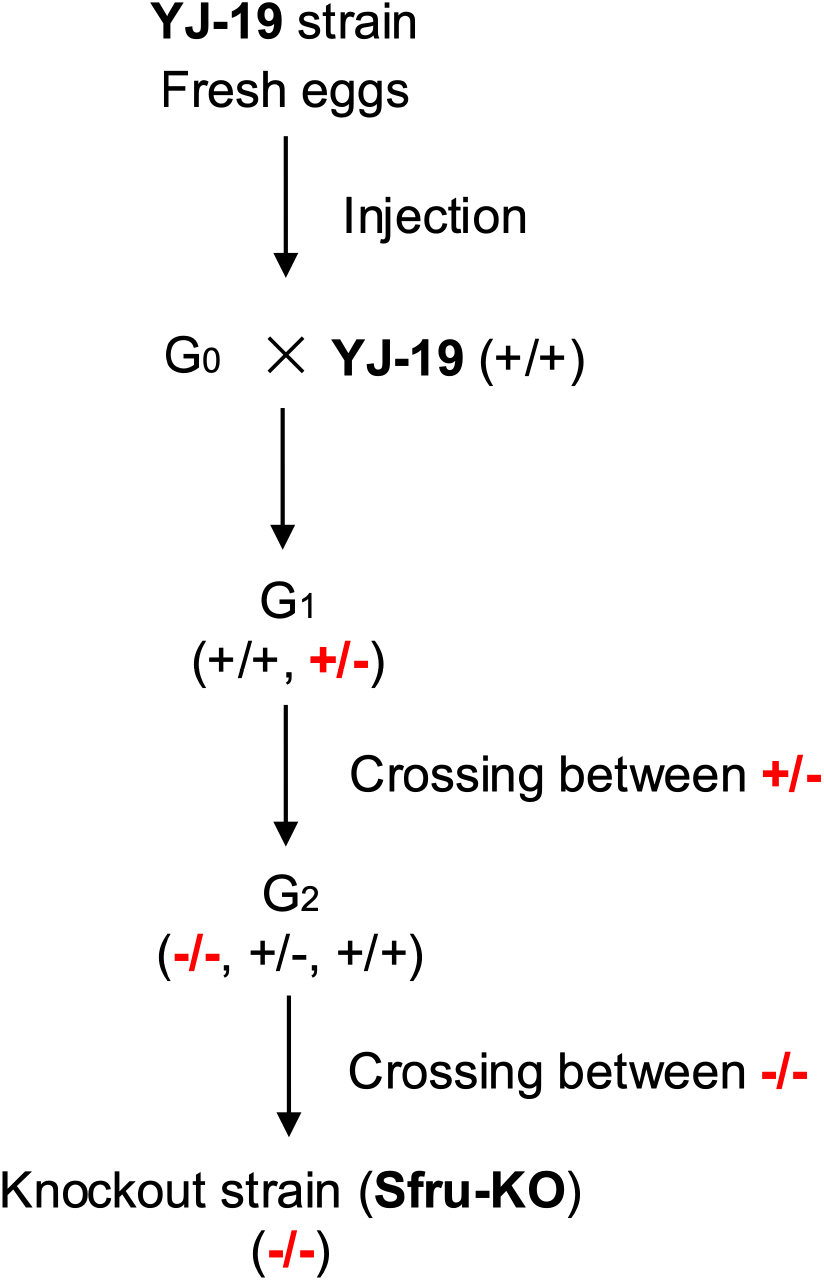
Diagram detailing the crossing and selection programs for establishing a homozygous knockout strain of *Spodoptera frugiperda*. (+/+) means the wild type, (-/-) means mutant homozygote, and (+/-) means mutant heterozygote.

**Figure 3.**
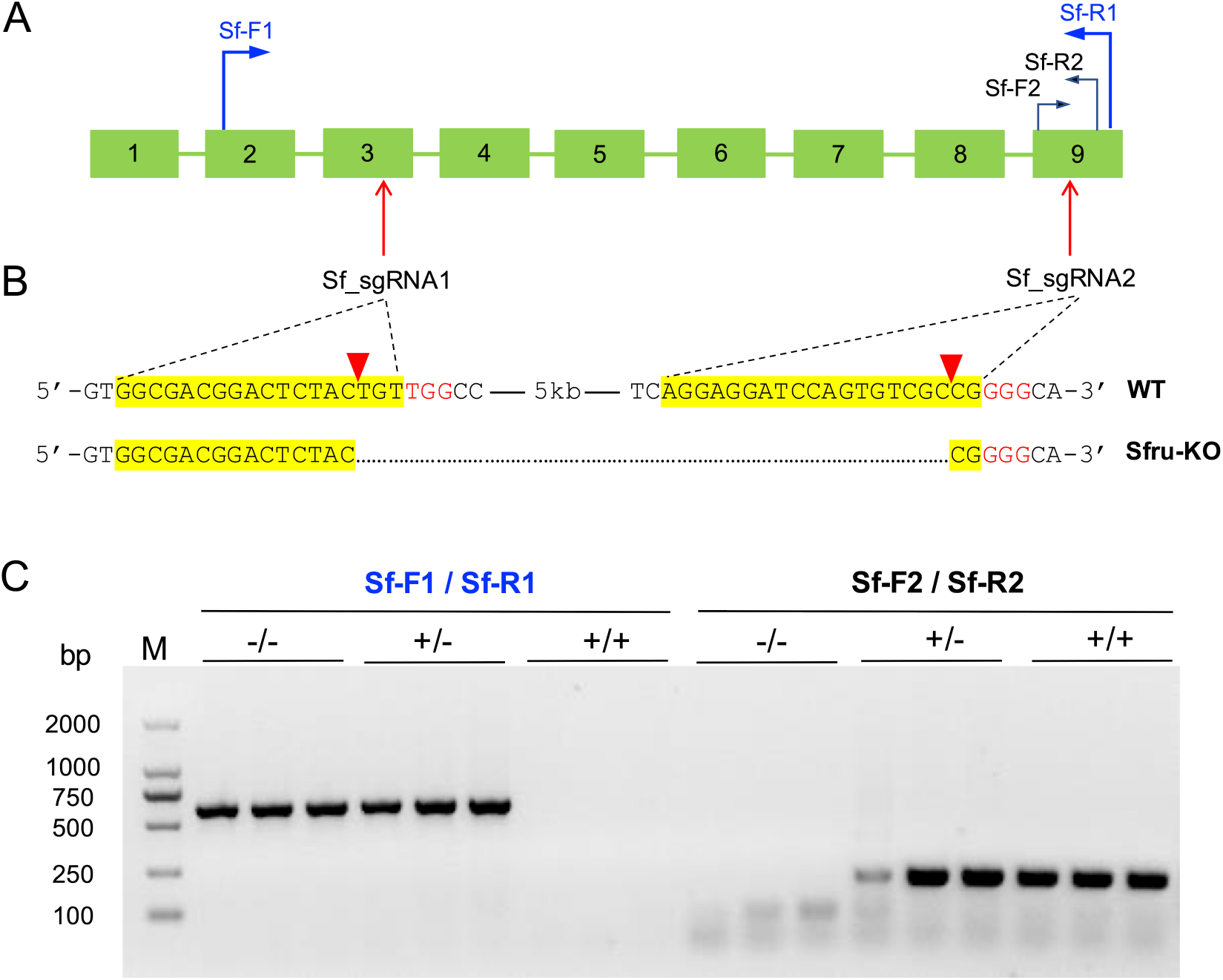
CRISPR/Cas9-mediated knockout of *Spodoptera frugiperda VipR1* (*SfVipR1*). (**A**) The genomic structure of *SfVipR1*, showing positions of two sgRNAs for gene editing and two primer pairs for allele-specific PCR detection. (**B**) Partial sequences of the wild-type and large fragment deletion alleles of *SfVipR1*. The targeting sequences of the two sgRNAs are highlighted in yellow, and the PAM sequences are shown in red. A red triangle indicates the cleavage site. The dotted line indicates the deleted base pairs. (**C**) Genotyping of individual *S. frugiperda* based on banding patterns from allele-specific PCR products.

### 3.4 Responses of the SfVipR1 knockout strain to Bt proteins Vip3Aa and Cry1Fa

Previous studies demonstrated that *HaVipR1* plays a critical role in the toxicity of Vip3Aa in *Helicoverpa armigera*. To determine whether *SfVipR1* has the same effects on Vip3Aa toxicity against *S. frugiperda*, the changes of the susceptibility to Vip3Aa were checked between the wild type and knockout strains.

Bioassay results (**Table 2**) showed that the SfVipR1 knockout strain (Sfru-KO) obtained >1875-fold resistance to Vip3Aa compared to the wild type YJ-19 strain of *S. frugiperda*. As expected, the knockout strain did not have a significant difference in susceptibility to Bt Cry protein Cry1Fa in comparison with the background YJ-19 strain.

**Table 2.**
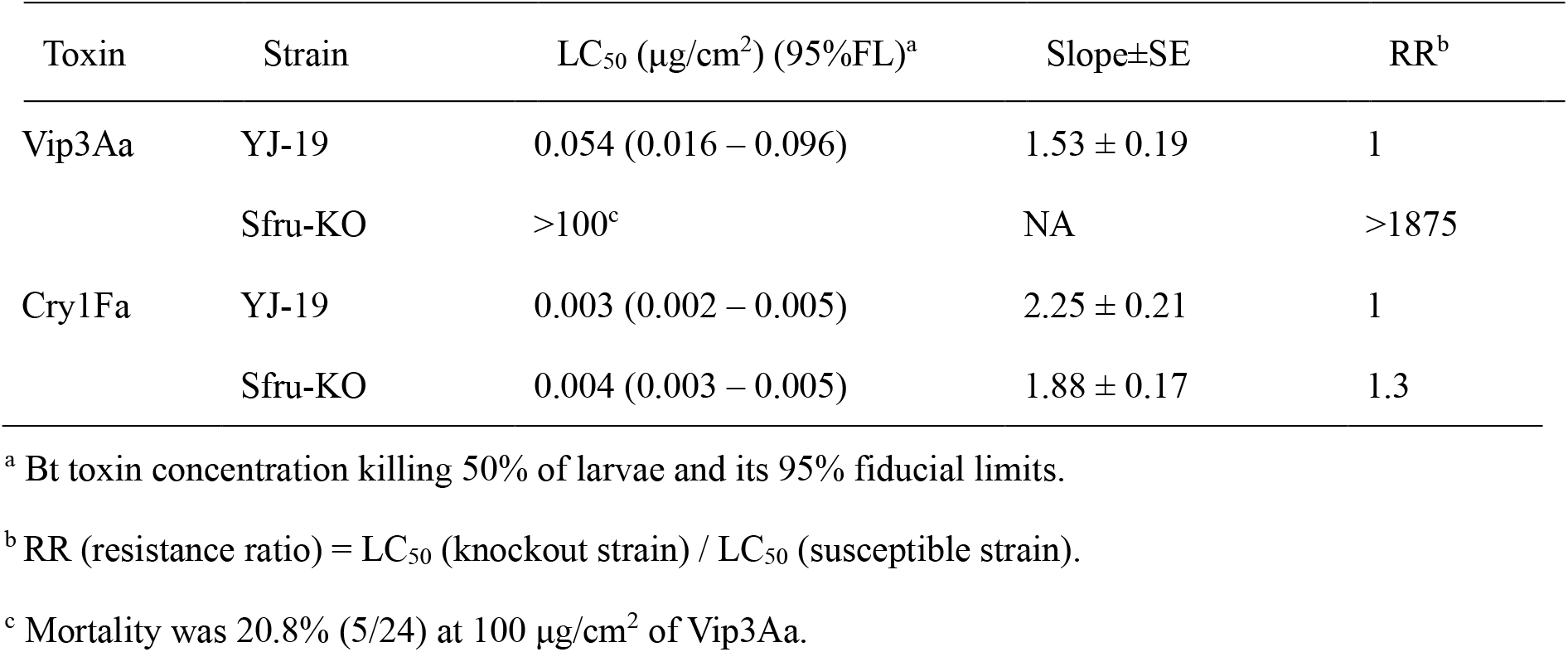
Susceptibility to Bt toxins Vip3Aa and Cry1Fa of the susceptible strains (YJ-19) and the knockout strain (Sfru-KO) of *S. frugiperda*.

| Toxin | Strain | LC<sub>50</sub> (µg/cm<sup>2</sup>) (95%FL)<sup>a</sup> | Slope±SE | RR<sup>b</sup> |
| --- | --- | --- | --- | --- |
| Vip3Aa | YJ-19 | 0.054 (0.016 – 0.096) | 1.53 ± 0.19 | 1 |
|  | Sfru-KO | >100<sup>c</sup> | NA | >1875 |
| Cry1Fa | YJ-19 | 0.003 (0.002 – 0.005) | 2.25 ± 0.21 | 1 |
|  | Sfru-KO | 0.004 (0.003 – 0.005) | 1.88 ± 0.17 | 1.3 |

<sup>a</sup> Bt toxin concentration killing 50% of larvae and its 95% fiducial limits.
<sup>b</sup> RR (resistance ratio) = LC<sub>50</sub> (knockout strain) / LC<sub>50</sub> (susceptible strain).
<sup>c</sup> Mortality was 20.8% (5/24) at 100 µg/cm<sup>2</sup> of Vip3Aa.

### 3.5 Inheritance of resistance to Vip3Aa in Sfru-KO strain

Under a treatment with Vip3Aa discriminating concentration (2 μg/cm^2^), survival was 0% for YJ-19 and 97.9% for Sfru-KO (**Table 3**). At this concentration, survival was 0% for the reciprocal F_1_ progeny between Sfru-KO and YJ-19 strains of *S. frugiperda*, and the value of the dominance parameter *h* is 0 (**Table 3**), indicating completely recessive inheritance. These results indicate inheritance of resistance to Vip3Aa was autosomal and recessive in the Sfru- KO strain of *S. frugiperda*.

**Table 3.**
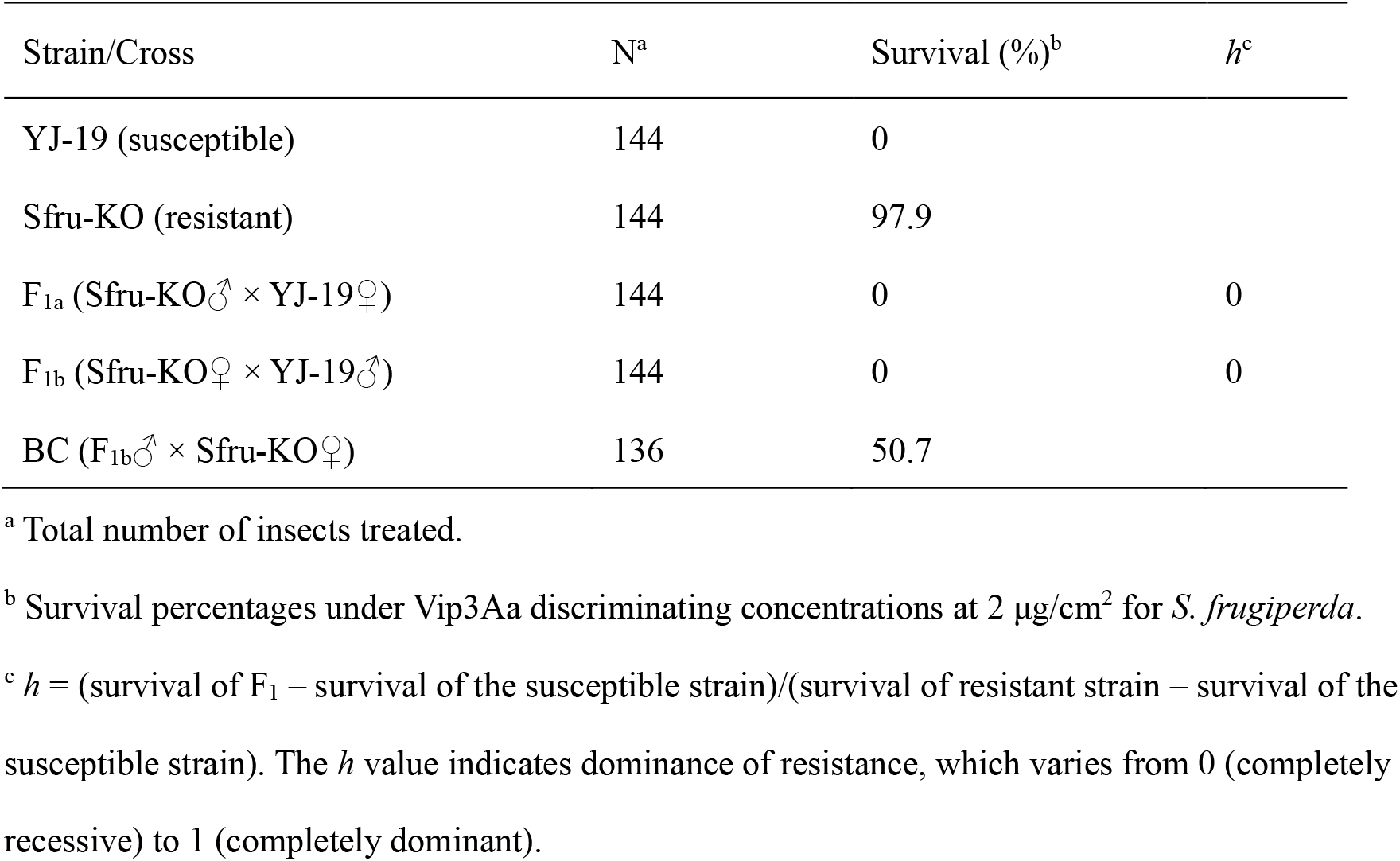
Inheritance of resistance to Vip3Aa in *S. frugiperda* Sfru-KO strain.

### 3.6 Genetic linkage between the SfVipR1 gene mutations and resistance to Vip3Aa

For genetic linkage analysis between the *SfVipR1* deletion mutation and Vip3Aa resistance, the male adults of F_1_ progeny between Sfru-KO and YJ-19 were pooled and back crossed with female adults of the resistant Sfru-KO strain of *S. frugiperda*. A total of 136 newly hatched larvae of the backcross progeny were treated with 2 μg/cm^2^ of Vip3Aa, and 50.7% larvae (69/136) survived (**Table 3**). All 69 survivors were genotyped to be homozygous for the deletion mutation of *SfVipR1*. In contrast, 20 untreated individuals randomly selected from the backcross progeny contained 11 mutant homozygotes and 9 mutant heterozygotes, which is in agreement with the expected 1:1 ratio (Fisher’s exact test, *p* = 1). The result demonstrates that the SfVipR1 knockout is significantly linked with Vip3Aa resistance in the Sfru-KO strain of *S. frugiperda* (Fisher’s exact test, *p* < 0.0001).

## 4 DISCUSSION

Genetic mapping and transcriptomic analyses have identified that disruption or downregulation of the *HaVipR1* gene is associated with high-level resistance to Vip3Aa in each of the two field-derived Australian *H. armigera* strains. Subsequent gene editing tests confirmed that the knockout of *HaVipR1* in an Africa-derived SCD strain of *H. armigera* produced a high-level of resistance to Vip3Aa, which is comparable to that of the resistant Australian strains.^22^ In the present study, knockout of *SfVipR1* also conferred high-level resistance to Vip3Aa in *S. frugiperda*. Another two Vip3A resistance genes have recently been reported in laboratory-selected resistant strains of *S. frugiperda*, the transcription factor *SfMyb* ^20^ and the chitin- synthase gene *SfCHS2*.^21^ It suggests there are several major Vip3A resistance genes in lepidopteran pests.

Five resistant strains to Vip3Aa of *H. zea* were established through F_2_ screens from field populations collected from three states (Texas, Louisiana, and Mississippi) in the United States.^16,30^ These five strains of *H. zea* are all highly resistant to Vip3Aa, and the resistance is inherited as a single, recessive and autosomal trait.^31,32^ However, three distinct genetic loci underlying Vip3Aa resistance were revealed through genetic complementary tests among these five resistant strains of *H. zea*.^32^ Similarly, Vip3Aa resistance in two *S. frugiperda* strains isolated respectively from Louisiana and Texas with F_2_ screens is recessive, autosomal and monogenic.^33,34^ The genetic complementary tests indicated these two strains have different genetic basis.^34^ In Brazil, four *S. frugiperda* strains highly resistant to Vip3Aa were derived from F_2_ screens of field populations from three states, and Vip3Aa resistance in the four strains is autosomal, recessive, and shares a common genetic locus.^11,15^ The above results demonstrate a diverse genetic basis for Vip3Aa resistance, at least in the field populations of *H. zea* and *S. frugiperda*, which could pose challenges for resistance detection and management. Identifying the genetic mechanisms behind these field-derived resistant strains will enhance our understanding of the mode of action of, and resistance to, Vip3A proteins, and consequently improve the detection and management of resistance in these key pests targeted by Bt crops. It would be of great interest to determine whether *VipR1, Myb* or *CHS2* is involved in the Vip3Aa resistance in these field-derived resistant strains reported in the previous studies.

The insecticidal mechanism of Vip3 proteins is supposed to follow a general pattern similar to that of Cry proteins: after the proteins are ingested, they are activated by the proteases within the larval midgut, pass through the peritrophic matrix, bind to specific receptor(s) in the apical membrane of the midgut epithelial cells, insert into the membrane, and cause colloid-osmotic cell lysis.^19^ Interactions between the Cry/Vip proteins and specific receptors are considered one of the key determinants of insecticidal activity. ABC transporters and cadherins have been identified as key functional receptors for Cry1A proteins in various lepidopteran insects.^8,35,36^ The specific receptors for Vip3A proteins have yet to be identified in lepidopteran insects, although several putative receptors for Vip3Aa have been suggested.^8,19^ Disruption of HaVipR1 results in high-level and recessive resistance to Vip3Aa in both *H. armigera*^22^ and *S. frugiperda* (this study), indicating that VipR1 plays a crucial role in mediating the toxicity of Vip3Aa in these two lepidopteran species. However, little is known about the functions of VipR1 in insects, or its role in the mode of action of Vip3Aa. It will be necessary in the future to investigate and ascertain the exact role of VipR1 in mediating the toxicity of Vip3Aa to its target pests, including *H. armigera* and *S. frugiperda*.

Decreased toxin binding is often associated with high levels of resistance to Cry proteins in lepidopteran insects.^8,18^ However, reduced binding of Vip3Aa was not found in the resistant strains of *H. armigera*^5^, *Heliothis virescens*^37^ and *Mythimna separata*.^38^ A recent study has detected a 3.7-fold reduction in the binding affinity of Vip3Aa with larval midguts in a Vip3Aa- resistant strain of *H. zea*.^39^ Variations in binding assays and the insect species examined might account for the differences between the binding results from these studies.^5,37-39^ Disruption of *HaVipR1* is causally associated with Vip3Aa resistance in the SP85 strain of *H. armigera*,^22^ while no decreased binding was observed in this resistant strain.^5^ The *SfVipR1* knockout strain of *S. frugiperda* created in the current study serves as an ideal subject for investigating toxin binding and additional biochemical mechanisms of Vip3Aa resistance in the future.

In China, invasive *S. frugiperda* was first officially reported in Yunnan Province in January 2019,^40^ and subsequent whole-genome sequencing and susceptibility bioassays have indicated that geographical populations of *S. frugiperda* collected from China during 2019-2022 remain susceptible to both Cry1Fa and Vip3Aa.^41,42^ Since 2019, the Ministry of Agriculture and Rural Affairs of China has been issuing safety certificates for several Bt corn hybrids that produce Cry1 and/or Vip3Aa proteins, and these Bt corn hybrids will be approved soon for commercial planting in China to control key lepidopteran pests, including *S. frugiperda*.^43^ With Asia and Southeast Asia identified as biosecurity hotspots underpinning the successful invasion and establishments of *S. frugiperda*,^44^ and considering that practical resistance to Cry1Fa has become widespread in the Americas due to the extensive use of Bt corn, it is critical to implement a resistance management (IRM) strategy when planting Bt corn in China^45^ and other Asian/Southeast Asian regions. This IRM strategy should incorporate high-dose/refuge and pyramid approaches to ensure the long-term effectiveness of Bt corn in China^45,46^ and surrounding areas. Once Bt corn is commercially planted, it is also imperative to implement efficient resistance monitoring programs to facilitate adaptive resistance management. The Sfru-KO strain of *S. frugiperda*, generated in this study, can be harnessed in an F_1_ screen program to determine the frequencies of resistance alleles of the *SfVipR1* gene associated with Vip3Aa resistance in *S. frugiperda*.

## ACKNOWLEDGEMENTS

This work was funded by the National Key Research Development Program of China (No. 2019YFD0300103) and the Fundamental Research Funds for the Central Universities of China (KYZ201920).

## DATA AVAILABILITY STATEMENT

The data that support the findings of this study are available from the corresponding author upon reasonable request.

